# Triptolide ameliorates renal tubulointerstitial fibrosis through EZH2

**DOI:** 10.1101/2023.01.29.526092

**Authors:** Yanzhe Wang, Ying Jing, Di Huang, Ming Wu, Chaoyang Ye

## Abstract

Renal fibrosis is the final pathological pathway of various kidney disease in progression to the end stage of renal failure. Recent studies showed that the histone methyltransferase enhancer of zeste homolog 2 (EZH2) is an important epigenetic regulator of renal fibrosis by promoting epithelial-mesenchymal transition (EMT) and activation of TGF-β/Smad3 signaling pathway in fibrotic kidneys. Triptolide is component extracted from radix tripterygium wilfordii, which provides renal benefits to patients with renal diseases in China. Recently, triptolide was identified as an inhibitor of EZH2. Thus, we hypothesized that triptolide inhibits renal fibrosis through EZH2. In this study, we found that triptolide reduced the deposition of extracellular matrix in the kidney of unilateral ureteral obstruction (UUO) mice. The anti-fibrotic effect of triptolide was further confirmed in TGF-β stimulated HK2 cells, a human renal epithelial cell line. Moreover, treatment of triptolide blocked the up-regulation of EZH2 in UUO kidneys and reduced the expression of EZH2 in TGF-β stimulated HK2 cells. Down-regulation of EZH2 by triptolide was correlated with reduced expression of EMT markers and phosphorylation of Smad3 in UUO kidneys and TGF-β stimulated HK2 cells. Finally, we showed that inhibition of EZH2 by 3-DZNep attenuated the inhibitory effect of triptolide on the expression of extracellular matrix protein, EMT markers and activation of Smad3 in TGF-β stimulated HK2 cells.

In conclusion, triptolide inhibits renal tubulointerstitial fibrosis through EZH2 in obstructive kidneys.

## Introduction

Chronic kidney disease (CKD) is an important public health and the prevalence of CKD is more than 10% of the world’s population(1). While CKD progresses to the terminal stage, renal fibrosis is the common pathway and the final pathological manifestation of CKD (2). It is characterized by the excessive deposition of extracellular matrix (ECM) in the kidney, such as fibronectin (FN) and collagen I (Col-I) in interstitial areas (3).

Triptolide (TP) is a diterpenoid trioxide isolated from the medicinal plant *Tripterygium wilfordii Hook F* (TWHF) (4, 5). Pharmacological studies have shown that triptolide exhibits renal protection through multiple effects such as anti-fibrotic, anti-inflammatory, and immunosuppressive effects (6). Extracts of TWHF have been used in the treatment of nephritis, minimal change disease, and membranous nephropathy in humans(7). Some animal model studies have shown that TP can decrease interstitial collagen deposition, inhibit renal interstitial fibroblast activation and attenuate renal tubular epithelial-mesenchymal transition(EMT) (8). However, the mechanisms underlying triptolide mediated renal protection is not completely known.

The histone methyltransferase enhancer of zeste homolog 2 (EZH2), as the catalytic part of polycomb repressive complex 2, is involved in fibrogenesis(9). It has been reported that blocking EZH2 activation with 3-deazaneplanocin A (3-DZNeP), a carbocyclic analog of adenosine, inhibits the activation of renal interstitial fibroblasts and attenuates the development of renal fibrosis (10). Our previous study also showed that inhibition of EZH2 by 3-DZNeP attenuated the anti-fibrotic effect of emodin in UUO kidneys and human renal epithelial cells (HK2) cells (11). A recent study showed that EZH2 is the direct target of triptolide(12).

In this study, we aimed to study the therapeutic effect of triptolide on renal tubulointerstitial fibrosis in a mouse model, and investigated the hypothesis that EZH2 mediates the anti-fibrotic effect of triptolide in renal fibrosis.

## Results

### Triptolide inhibits attenuates tubulointerstitial fibrosis in obstructive mouse kidneys and renal cells

The effect of triptolide on renal tubulointerstitial fibrosis was evaluated by using unilateral ureteral obstruction (UUO) mice. Massive interstitial collagen deposition was observed in UUO kidneys at 10 days after the operation as shown by Masson staining, and treatment with triptolide attenuated collagen deposition (Figure 1A-1B). Increased ECM markers such as collagen-I and fibronectin were shown in UUO kidneys by Western blotting and triptolide reduced the expression of these two ECM markers (Figure 1C-1D).

**Figure 1.**
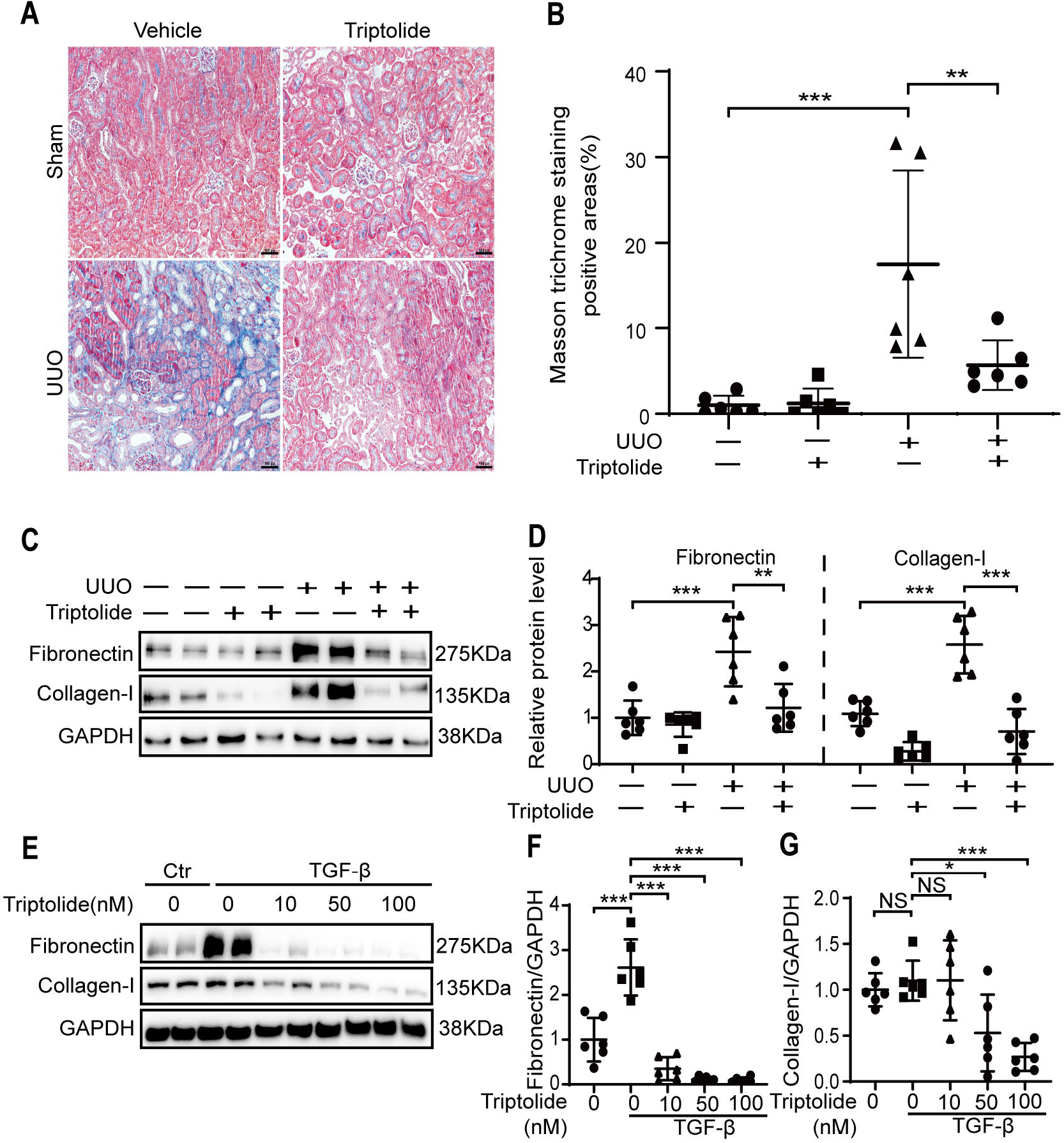
Triptolide inhibits attenuates tubulointerstitial fibrosis in obstructive mouse kidneys and renal cells. Sham or unilateral ureter obstruction (UUO) operation was performed on wildtype c57 mice, followed by 10 days of treatment with DMSO or triptolide. **A and B**. Renal fibrosis was assessed by Masson’s trichrome staining and then quantified. Bars◻=◻100 μm. **C and D**. The expression of fibronectin and collagen I were analyzed by Western blotting and then quantified. **E-G**. HK2 human renal epithelial cells were starved overnight and followed by stimulation with TGF-β and treatment with various concentration of triptolide for 48h. Cell lysates were extracted. The expression of fibronectin and collagen I were analyzed by Western blotting and then quantified. Data represents mean◻±◻SD. One representative result of at least three independent experiments is shown. NS represents not significant. \**p*◻<◻0.05. \*\**p*◻<◻0.01. \*\*\**p*◻<◻0.001.

The anti-fibrotic effect of triptolide was further confirmed *in vitro* by using TGF-β stimulated human renal epithelial (HK2) cells. Triptolide dose-dependently inhibited the expression of fibronectin and collagen-I in TGF-β stimulated HK2 cells from 10 nM to 100 nM (Figure 1E-1G).

### Triptolide inhibits the expression of EZH2 in *in vitro* and *in vivo* model of renal fibrosis

Next, we studied the effect of triptolide on EZH2 in renal fibrosis. As shown in Figure 2A and 2B, EZH2 was up-regulated in UUO kidneys as compared with that in sham kidneys, and treatment with triptolide reduced the expression of EZH2 in UUO kidneys.

**Figure 2.**
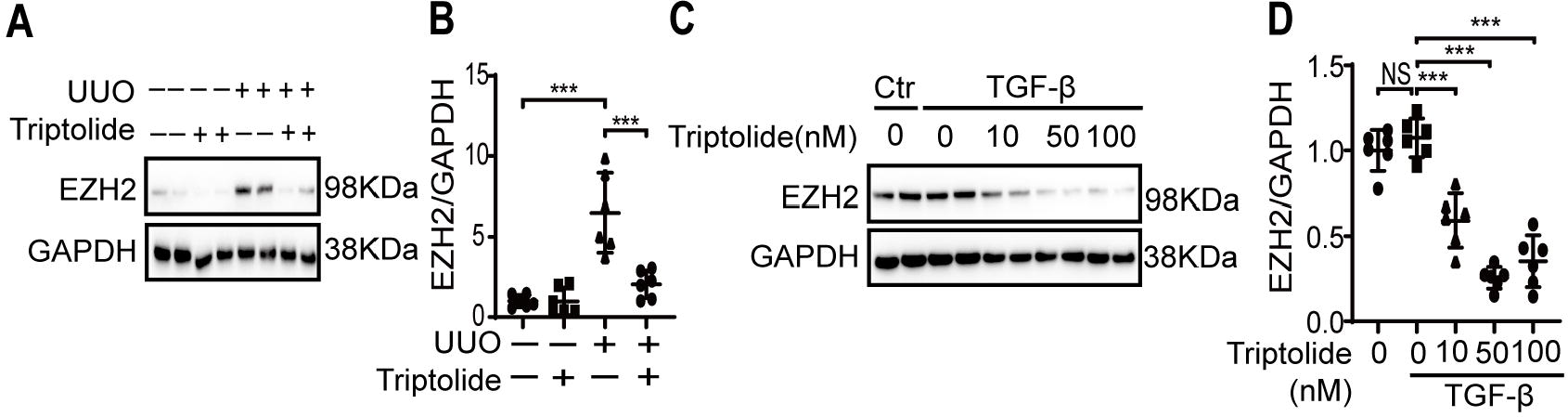
Triptolide inhibits the expression of EZH2 in *in vitro* and *in vivo* model of renal fibrosis. **A and B**. The expression of EZH2 in sham or UUO kidneys was analyzed by Western blotting and further quantified. **C and D**. HK2 human renal epithelial cells were starved overnight, then stimulated with TGF-β and treated with various concentrations of triptolide for 48h. The expression of EZH2 was analyzed by Western blotting and further quantified. One representative of at least three independent experiments is shown. Data represents the mean◻±◻SD. NS represents not significant. \*\*\**p*◻<◻0.001.

We further showed that triptolide inhibited the expression of EZH2 in TGF-β stimulated HK2 cells from 10 nM to 100 nM (Figure 2C-2D).

### Triptolide reduces the expression of EMT markers and phosphorylation of Smad3 in fibrotic mouse kidneys and renal cells

Since EZH2 promotes EMT and activates the TGF-β/Smad3 signaling pathway in fibrotic kidneys (10, 13), we next investigated the effect of triptolide on EMT and Smad3 phosphorylation in fibrotic kidneys.

As shown in Figure 3A to 3D, three EMT markers vimentin, a-SMA and Snail were increased in UUO kidneys, which were reduced by the treatment of triptolide. Similarly, the expression of EMT markers (N-cadherin, vimentin and Snail) were dose-dependently inhibited by triptolide in TGF-β stimulated HK2 cells from 10 nM to 100 nM (Figure 3E-3G).

**Figure 3.**
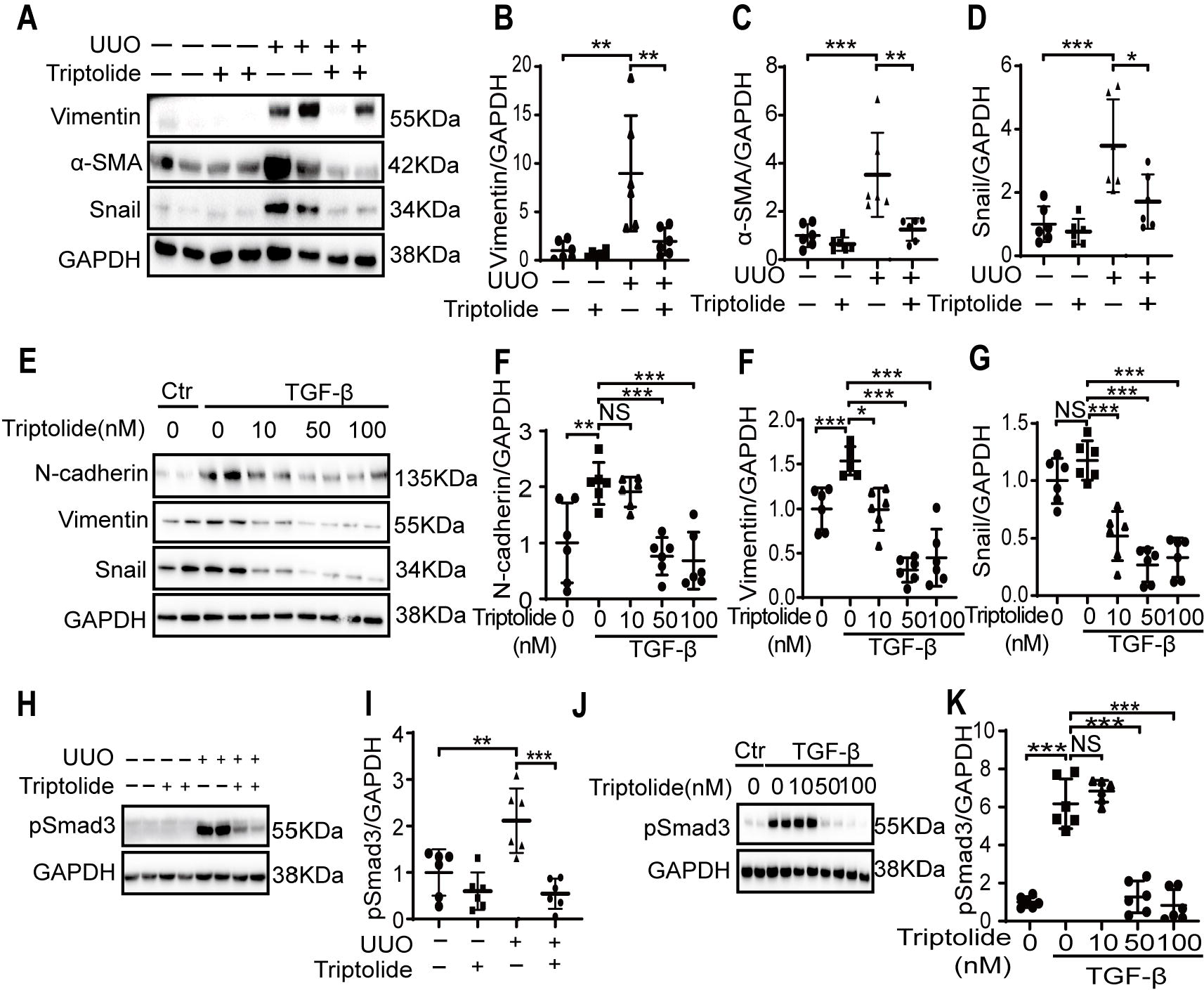
Triptolide reduces the expression of EMT markers and phosphorylation of Smad3 in fibrotic mouse kidneys and renal cells. **A-D**. In sham or UUO kidneys, the expression of vimentin,α-SMA and snail were analyzed by Western blotting and then quantified. **E-G**. HK2 human renal epithelial cells were starved overnight, then stimulated with TGF-β and treated with various concentrations of triptolide for 48h. The expression of N-cadherin, vimentin and snail were analyzed by Western blotting and then quantified. **H and I**. The expression of phosphorylated Smad3 (pSmad3) in sham or UUO kidneys was analyzed by Western blotting and then quantified. **J and K**. The expression of phosphorylated Smad3 (pSmad3) in HK2 human renal epithelial cells was analyzed by Western blotting and then quantified. One representative of at least three independent experiments is shown. Data represents the mean◻±◻SD. NS represents not significant.\**p*◻<◻0.05. \*\**p*◻<◻0.01. \*\*\**p*◻<◻0.001.

Activation of Smad3 was shown in UUO kidneys or TGF-β stimulated HK2 cells as compared with their controls (Figure 3H-3K). Triptolide inhibited the phosphorylation of Smad3 (pSmad3) in UUO and TGF-β stimulated HK2 cells (Figure 3H-3K).

### Triptolide inhibits renal fibrosis and EMT through EZH2

To determine whether EZH2 is involved in the anti-fibrotic effect of triptolide, we used EZH2 specific inhibitor 3-DZNep to inhibit the expression of EZH2 in TGF-β stimulated HK2 cells (Figure 4A-4B). Deletion of EZH2 by 3-DZNep reduced the expression of fibronectin and collagen-I in TGF-β stimulated HK2 cells and diminished or blocked the inhibitory effect of triptolide on fibronectin and collagen-I expression (Figure 4C-4E). We further showed that 3-DZNep diminished the inhibitory effect of triptolide on EMT markers (N-cadherin, vimentin and Snail) and activation of Smad3 (Figure 4F-4G). These data suggested that triptolide inhibits renal fibrosis through EZH2.

**Figure 4.**
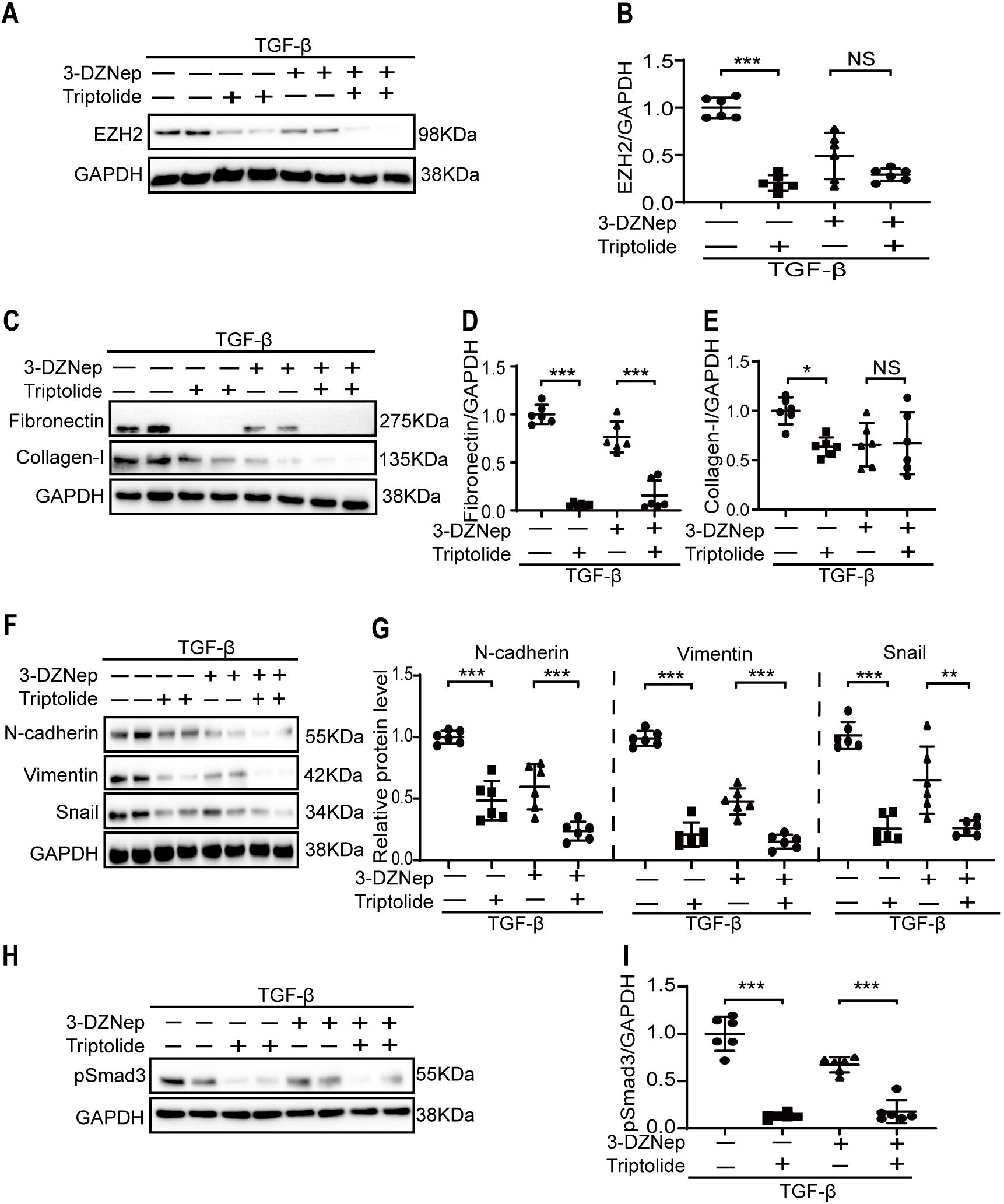
Triptolide inhibits renal fibrosis and EMT through EZH2. HK2 human renal epithelial cells were starved overnight and followed by stimulation with TGF-β and treatment with 20 μM of 3-DZNeP and 100nM of triptolide for 48h. Cell lysates were extracted. **A and B**. The expression of EZH2 was analyzed by Western blotting and further quantified. **C-E**. The expression of fibronectin and collagen I were analyzed by Western blotting and then quantified. **F and G**. The expression of N-cadherin, vimentin and snail were analyzed by Western blotting and then quantified. **H and I**. The expression of phosphorylated Smad3 (pSmad3) was analyzed by Western blotting and then quantified. One representative of at least three independent experiments is shown. Data represents the mean◻±◻SD. NS represents not significant.\**p*◻<◻0.05. \*\**p*◻<◻0.01. \*\*\**p*◻<◻0.001.

## Discussion

Triptolide, a natural compound originally extracted from the traditional Chinese medicine Tripterygium wilfordii, has been used to treat chronic kidney diseases. The anti-fibrotic effects have been reported in some of the animal models (14, 15). To determine the effect of triptolide on renal tubulointerstitial fibrosis, we established the UUO mice model of renal fibrosis in this study. We showed that 10 days treatment with triptolide reduced collagen deposition by Masson staining in UUO kidneys. Western blotting analysis showed the down-regulation of pro-fibrotic markers such as fibronectin and collagen-I by triptolide treatment in the UUO mouse models. We also showed that triptolide inhibited the expression of EMT markers and pSmad3. The direct anti-fibrotic effect of triptolide on renal cells was further studied using HK2 cells. We showed that triptolide dose-dependently reduced the expression of ECM protein, several EMT markers, and phosphorylation of Smad3. Thus, we conclude that triptolide inhibits renal tubulointerstitial fibrosis.

EZH2, a histone-lysine N-methyltransferase enzyme, has been identified as an important therapeutic target to treat renal tubulointerstitial fibrosis (16). EZH2 is up-regulated in several animal models of renal fibrosis and pharmaceutically inhibition of EZH2 ameliorates renal tubulointerstitial fibrosis (17). Recent study showed that EZH2 is a direct target of triptolide and thus we hypothesized that triptolide inhibits renal fibrosis by targeting EZH2. In this study, we showed that triptolide inhibited the expression of EZH2 in the UUO model which was correlated with reduced renal fibrosis. *In vitro*, triptolide dose-dependently inhibited the expression of EZH2 which is tightly correlated with the down-regulation of pro-fibrotic markers. Inhibition of EZH2 by 3-DZNep attenuated the anti-fibrotic effect.

We conclude that T triptolide inhibits renal interstitial fibrosis through the inhibition of EZH2.

## Materials and Methods

### Animal studies

Wide type male C57BL/6 mice with body weight between 20 to 25g were purchased from Shanghai SLAC Laboratory Animal Co., Ltd and housed in a SPF grade animal facility in the Shanghai University of Traditional Chinese Medicine under the local regulations. The animal experimentation ethics committee of Shanghai University of Traditional Chinese Medicine has approved the animal experiments (PZSHUTCM18111601).

UUO operation was performed through twice ligation of the left ureter with 4-0 nylon sutures. WT animals were sacrificed at day 3, 7 or 14 in a time-course experiment (n=5 for each time point). For interventional study, WT mice were randomly divided into four groups: 1) Sham/Vehicle (n=6), 2) Sham/triptolide (n=7), 3) UUO/ Vehicle (n=7), and 4) UUO/triptolide (n=7) group. The stock of triptolide (Topscience, T2179, Shanghai, China) was dissolved in DMSO and further diluted in normal saline as working solution. Sham or UUO mice were treated with vehicle or 0.25 mg/kg triptolide daily by intraperitoneal (i.p.) injection for 10 days starting from day 0. Mice were sacrificed at day 10. Kidney samples were obtained for Western blotting analysis or histological examinations.

### Cell culture

HK2 renal proximal tubular epithelial cells were obtained from the Cell Bank of Shanghai Institute of Biological Sciences (Chinese Academy of Science). HK2 cells were seeded in 6-well plate to 40-50% confluence, which were starved overnight with DMEM/F12 medium containing 0.5% fetal bovine serum. On the next day, fresh medium containing 0.5% fetal bovine serum was changed, and then cells were exposed to 2.5 ng/ml TGF-β (Peprotech, Rocky Hill, NJ, USA) for 48h in the presence of various concentration of triptolide.

### Masson’s trichrome

Mouse kidneys were sliced, fixed, and embedded in paraffin, and cut into four-μm-thick sections. The paraffin-embedded kidney section was stained with hematoxylin, and then with ponceau red liquid dye acid complex, which was followed by incubation with phosphomolybdic acid solution. Finally, the tissue was stained with aniline blue liquid and acetic acid. Images were captured by using a microscope (Nikon 80i, Tokyo, Japan).

### Western blotting analysis

Cell or kidney protein was extracted by using lysis buffer bought from Beyotime Biotech (Nantong, China). BCA Protein Assay Kit (P0012S, Beyotime Biotech, Nantong, China) was used to determine the protein concentration. Protein samples were dissolved in 5x SDS-PAGE loading buffer (P0015L, Beyotime Biotech, Nantong, China), which were further subjected to SDS-PAGE gel electrophoresis. After electrophoresis, proteins were electro-transferred to a polyvinylidene difluoride membrane (Merck Millipore, Darmstadt, Germany). Unspecific binding on the PVDF membrane was blocked by incubation with the blocking buffer (5% non-fat milk, 20mM Tris-HCl, 150mM NaCl, PH=8.0, 0.01%Tween 20) for 1 hour at room temperature, which was followed by incubation with anti-fibronectin (1:1000, ab23750, Abcam), anti-Collagen I (1:500, sc-293182, Santa Cruz), anti-α-SMA (1:1000, ET1607-53, HUABIO), anti-EZH2 (1:1000, 5246S, Cell Signaling Technology) antibody, anti-N-cadherin (1:1000, sc-59887, Santa Cruz), anti-Vimentin (1:1000, R1308-6, HUABIO), anti-Snail(1:1000, A11794, ABclonal), anti-pSmad3 (1:1000, ET1609-41, HUABIO), anti-GAPDH (1:5000, 60004-1-lg, Proteintech) antibodies overnight at 4◻. Binding of the primary antibody was detected by an enhanced chemiluminescence method (SuperSignal™ West Femto, 34094, Thermo Fisher Scientific) using horseradish peroxidase-conjugated secondary antibodies (goat anti-rabbit IgG, 1:1000, A0208, Beyotime or goat anti-mouse IgG, 1:1000, A0216, Beyotime).

### Statistical analysis

Results were presented as mean ± SD. Differences among multiple groups were analyzed by one-way analysis of variance (ANOVA) and comparison between two groups was performed by unpaired student t-test by using GraphPad Prism version 8.0.0 for Windows (GraphPad Software, San Diego, California USA). A P value of lower than 0.05 was considered statistically significant.

## Author Contributions

CY funded the project. MW, YJ and CY conceived this project. MW coordinated the study and wrote the paper. YW conducted the *in vitro* experiments. MW, YW, YJ and DH performed the animal experiments. YW performed and analyzed the Western blotting. All authors reviewed the results and approved the final version of the manuscript.

